# Hexokinase 2 is dispensable for photoreceptor development but is required for survival during aging and outer retinal stress

**DOI:** 10.1101/782607

**Authors:** Eric Weh, Zuzanna Lutrzykowska, Andrew Smith, Heather Hager, Mercy Pawar, Thomas Wubben, Cagri G. Besirli

**Author notes:** Corresponding Author, please direct all correspondence to: Cagri Besirli, 1000 Wall St., Ann Arbor, MI 48105.

## Abstract

Photoreceptor death is the ultimate cause of vision loss in many retinal degenerative conditions. Identifying novel therapeutic avenues for prolonging photoreceptor health and function has the potential to improve vision and quality of life for patients suffering from degenerative retinal disorders. Photoreceptors are metabolically unique among other neurons in that they process the majority of their glucose via aerobic glycolysis. One of the main regulators of aerobic glycolysis is hexokinase 2 (HK2). Beyond its enzymatic function of phosphorylating glucose to glucose-6-phosphate, HK2 has additional non-enzymatic roles, including the regulation of apoptotic signaling via AKT signaling. Determining the role of HK2 in photoreceptor homeostasis may identify novel signaling pathways that can be targeted with neuroprotective agents to boost photoreceptor survival during metabolic stress. Here we show that following experimental retinal detachment, p-AKT is upregulated and HK2 translocates to mitochondria. Inhibition of AKT phosphorylation in 661W photoreceptor-like cells results in translocation of mitochondrial HK2 to the cytoplasm, increased caspase activity, and decreased cell viability. Rod-photoreceptors lacking HK2 upregulate HK1 and appear to develop normally. Interestingly, we found that HK2-deficient photoreceptors are more susceptible to acute nutrient deprivation in the experimental retinal detachment model. Additionally, HK2 appears to be important for preserving photoreceptors during aging. We show that retinal glucose metabolism is largely unchanged after HK2 deletion, suggesting that the non-enzymatic role of HK2 is important for maintaining photoreceptor health. These results suggest that HK2 expression is critical for preserving photoreceptors during acute nutrient stress and aging. More specifically, p-AKT mediated translocation of HK2 to the mitochondrial surface may be critical for protecting photoreceptors from acute and chronic stress.

## Introduction

Nearly 10 million individuals in the US alone are affected with either diabetic retinopathy, age-related macular degeneration, or retinitis pigmentosa, and the number of affected adults is only predicted to increase due to the aging population of the United States.^1^ Although the pathological mechanisms underlying these various retinal degenerations are complex, the ultimate cause of blindness is the death of photoreceptor cells. Therefore, identifying a common mechanism for promoting photoreceptor cell survival agnostic of the upstream pathological signaling has immense potential for preserving vision in many types of retinal diseases.

Photoreceptors require a tremendous amount of macromolecules to replenish their outer segments, which are shed daily.^2^ In addition, photoreceptors seem to possess little reserve capacity for energy production and are extremely sensitive to metabolic dysfunction as evidenced by mutations in common metabolism genes leading to isolated inherited retinal degenerations. For example, the enzymes inosine monophosphate dehydrogenase 1 (IMPDH1), hexokinase 1 (HK1), and isocitrate dehydrogenase 3 beta (IDH3B) are expressed throughout most tissues in the body; however, mutations affecting the genes encoding for these proteins result in isolated retinal degeneration.^3–5^ These data suggest that the metabolism of photoreceptors is tightly regulated and small perturbations may lead to deficiencies which cannot be compensated for, resulting in cell death.^6^ Therefore, identifying pathways important for cell survival during metabolic stress may prove useful for delaying or preventing photoreceptor cell death in many types of retinal degenerations.^7–10^ These and other data have led to a hypothesis suggesting that disruption of metabolism and nutrient availability may be a unifying mechanism for photoreceptor cell death in many degenerative retinal diseases.^10,11^

Photoreceptors utilize a type of metabolism unique among all other terminally differentiated neurons, called aerobic glycolysis or the Warburg effect, to process the majority of glucose that enters the cell.^12^ Photoreceptors may employ this unique metabolic adaptation to meet their high demand for energy and biosynthetic intermediates.^13^ Aerobic glycolysis tightly regulates substrate utilization, either allowing the shuttling of glycolytic intermediates to other biosynthetic pathways for nucleotide, amino acid, and lipid synthesis or to energy production via the oxidative phosphorylation pathway, depending on the physiologic needs of the cell.^14^ Three enzymes are key for performing aerobic glycolysis: hexokinase 2 (HK2), pyruvate kinase muscle isozyme 2 (PKM2) and lactate dehydrogenase A (LDHA). We have recently shown that genetically switching PKM2 for PKM1 protects photoreceptors from acute outer retinal stress induced by experimental retinal detachment.^7^ Interestingly, HK2 has been shown to play a role in mediating apoptotic cell processes after nutrient stress.^15,16^ This identifies HK2 as another attractive target for developing novel neuroprotective interventions in the retina.

Hexokinase 2 (HK2) is one of four main isozymes responsible for the first step in glycolysis. HK2 normally resides on the mitochondria, bound to voltage dependent anion channel (VDAC), where it receives preferential access to ATP for the phosphorylation of glucose to glucose-6-phosphate. While bound to VDAC, HK2 can also prevent the binding of pro-apoptotic factors to the mitochondria and the opening of the mitochondria permeability transition pore (mPTP), which blocks the release of cytochrome c.^16,17^ Protein kinase B (AKT) has been shown to modulate the ability of HK2 to bind to VDAC through direct phosphorylation.^18,19^

In this report, we show that HK2 preferentially localizes to the mitochondrial enriched fraction after outer retinal stress as produced by experimental retinal detachment. Disrupting the association between HK2 and mitochondria can be partially blocked by inhibiting AKT phosphorylation *in vitro*, which leads to decreased cell viability and increased caspase activity. We also demonstrate that rod photoreceptor-specific, *Hk2* conditional knockout (cKO) mice are more susceptible to acute outer retinal metabolic stress, suggesting an anti-apoptotic role for HK2 during metabolic stress. Additionally, we show that the loss of *Hk2* in rod photoreceptors does not reprogram metabolism to primarily oxidative phosphorylation. Finally, the rod photoreceptor-specific, *Hk2* cKO mice show significant outer retinal thinning and photoreceptor loss at advanced age. Collectively, these findings indicate that HK2 is critical for regulating photoreceptor survival during acute metabolic stress and normal aging.

## Results

### HK2 localizes to mitochondria following retinal detachment

One of the non-enzymatic roles of HK2 is to inhibit apoptosis by preventing opening of the mPTP.^15,16,18^ This non-enzymatic function can only be done while HK2 is physically bound to VDAC on the outer mitochondrial membrane. AKT has been shown to phosphorylate HK2, which promotes binding to VDAC.^18^ In order to determine if HK2 binding to VDAC is important for photoreceptor protection after retinal detachment, the levels of HK2 and the ratio of p-AKT/total AKT were assessed following experimental retinal detachment (RD) in rats (Fig. 1). Three and seven days following RD, total HK2 protein expression assessed by Western blot analysis was decreased significantly (Fig. 1a). Total AKT expression was unchanged, but p-AKT (S473) and the ratio of p-AKT/total AKT was significantly increased (Fig. 1b). To determine if this increase in p-AKT is associated with changes in HK2 sub-cellular localization, rat retinas were detached and harvested at 1-, 3- and 7-days post-RD. After fractionation, HK2 was found to be enriched in the post-cytosolic, mitochondrial enriched fraction (hereafter referred to as “mitochondrial fraction”) 3- and 7-days after retinal detachment (Fig. 1c&d), suggesting that increased p-AKT may be enhancing HK2 association with mitochondria.

**Figure 1.**
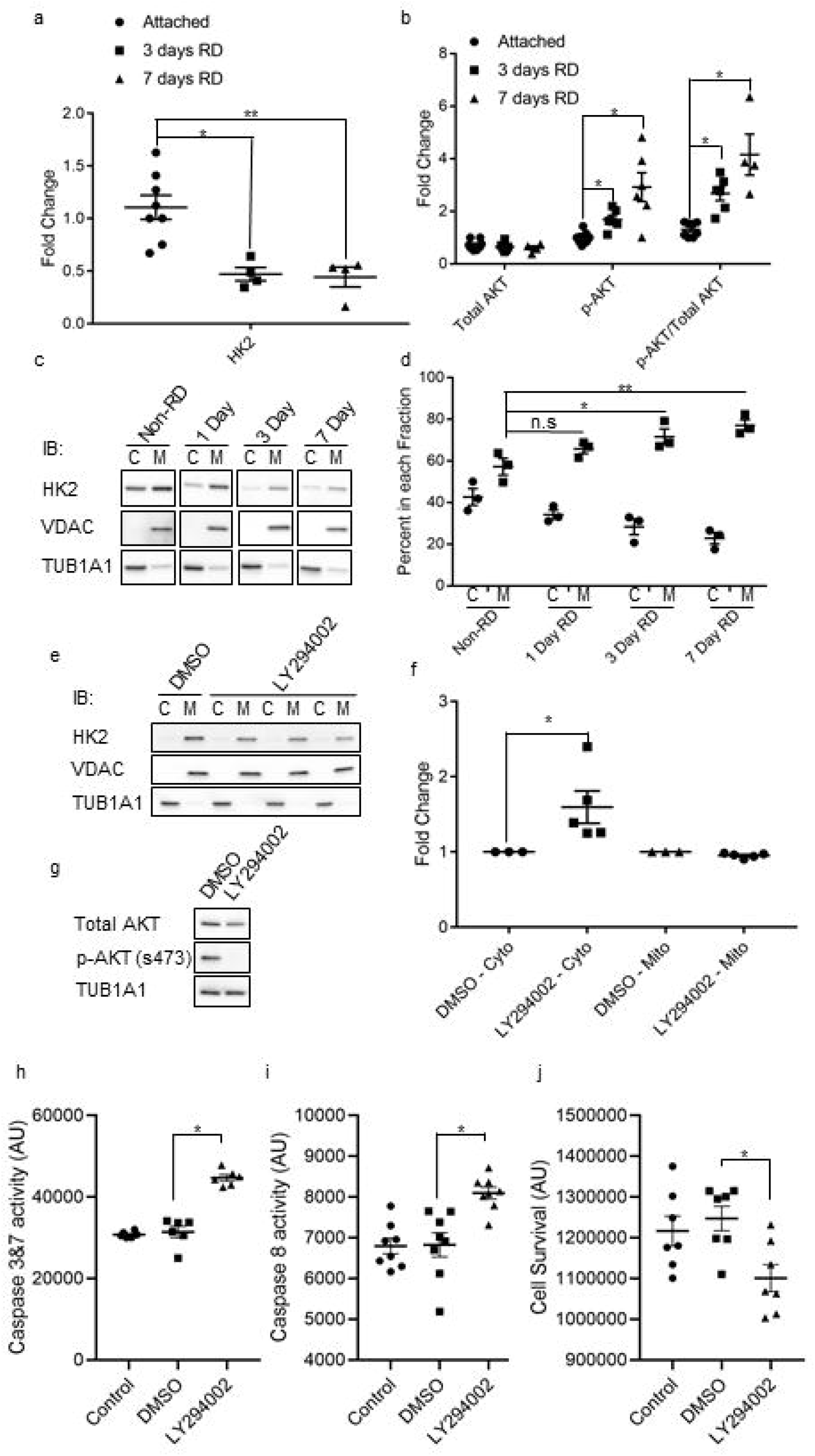
HK2 is differentially regulated after retinal detachment. (a) Total HK2 levels are significantly decreased 3- and 7-days post retinal detachment as assayed by Western blot. (b) Total AKT levels are unchanged after retinal detachment while p-AKT (S473) levels are significantly increased as assayed by Western blot. (c) Representative Western blots of fractionated rat retinas. VDAC was used as a mitochondrial fraction marker, TUB1A1 (α-tubulin) was used as a cytosolic fraction marker. (d) Percentage of HK2 signal in each fraction. HK2 is significantly enriched in the mitochondrial fraction 3- and 7-days after retinal detachment. (e) HK2 localization after 1.5 hours of treatment with 50μM LY294002 as assayed by Western blot. VDAC was used as a mitochondrial fraction marker, TUB1A1 (α-tubulin) was used as a cytosolic fraction marker. (f) quantification of data from e. (g) Whole cell lysate showing absence of p-AKT after 1.5 hours of 50μM LY294002 treatment as assayed by Western blot. (h, i & j) Caspase 3/7 and 8 activation and cell viability after 6 hours of 50μM LY294002 treatment. C-cytosolic fraction, M – mitochondrial enriched fraction, N=3-9, mean + SEM *-p≤.05, **-p<.01

### LY294002 treatment induces HK2 translocation to the cytosol and caspase activation

To better define the relationship between AKT and HK2 localization, the 661W photoreceptor-like cell model was used.^20,21^ Nearly all of the HK2 present in 661W cell extracts localizes to the mitochondrial fraction (Fig. 1e). When treated with LY294002 for 1.5 hours, a significant enrichment of HK2 in the cytosolic fraction was observed (Fig. 1f) and phosphorylation of AKT is nearly abolished as demonstrated on Western blot (Fig. 1g). After 6 hours of exposure to LY294002, both caspase 3/7 and 8 were significantly activated (Fig. 1h&i) with a corresponding significant decrease in cell viability (Fig. 1j). These data agree with previous studies demonstrating a neuroprotective effect of activated AKT on photoreceptors and suggest that at least part of this neuroprotective effect may be mediated by the regulation of HK2 localization in the cell.^19,22,23^

### Deletion of *Hk2* from rod photoreceptors leads to *Hk1* upregulation

Since the previous data suggest that HK2 may be important for preserving photoreceptor health during apoptotic stress, a rod photoreceptor-specific, *Hk2* conditional knockout mouse model was constructed to study the effects of *Hk2* deficiency in photoreceptors. Mice harboring a floxed *Hk2* gene were crossed with Rho-Cre mice, where Cre-recombinase is expressed specifically in rod photoreceptors.^24,25^ Mice with both *Hk2* alleles present in photoreceptors (*Hk2^wt/wt^;Rho-Cre*^+^: WT), and animals lacking both *Hk2* alleles in photoreceptors (*Hk2^fl/fl^;Rho-Cre^+^*: cKO) were produced. Consistent with other reports, immunofluorescence of retinal sections from WT mice showed that HK2 is almost exclusively expressed in photoreceptors (Fig. 2a). In cKO animals, the HK2 signal was only present in cone photoreceptors as shown by cone arrestin (ARR3) staining (Fig. 2b).^26,27^ Additionally, WT retinas express low levels of HK1 in photoreceptor inner segments (Fig. 2c) with the vast majority of HK1 expressed in other cells within the retina. Immunofluorescent staining of retinas from cKO mice demonstrated that photoreceptors upregulate HK1, and this increased HK1 expression is predominantly localized to rod inner segments (Fig. 2c). Western blotting confirmed these results by showing that HK2 is nearly absent from the retina of cKO animals while total HK1 levels are statistically significantly increased (Fig. 2d and 2e).

**Figure 2.**
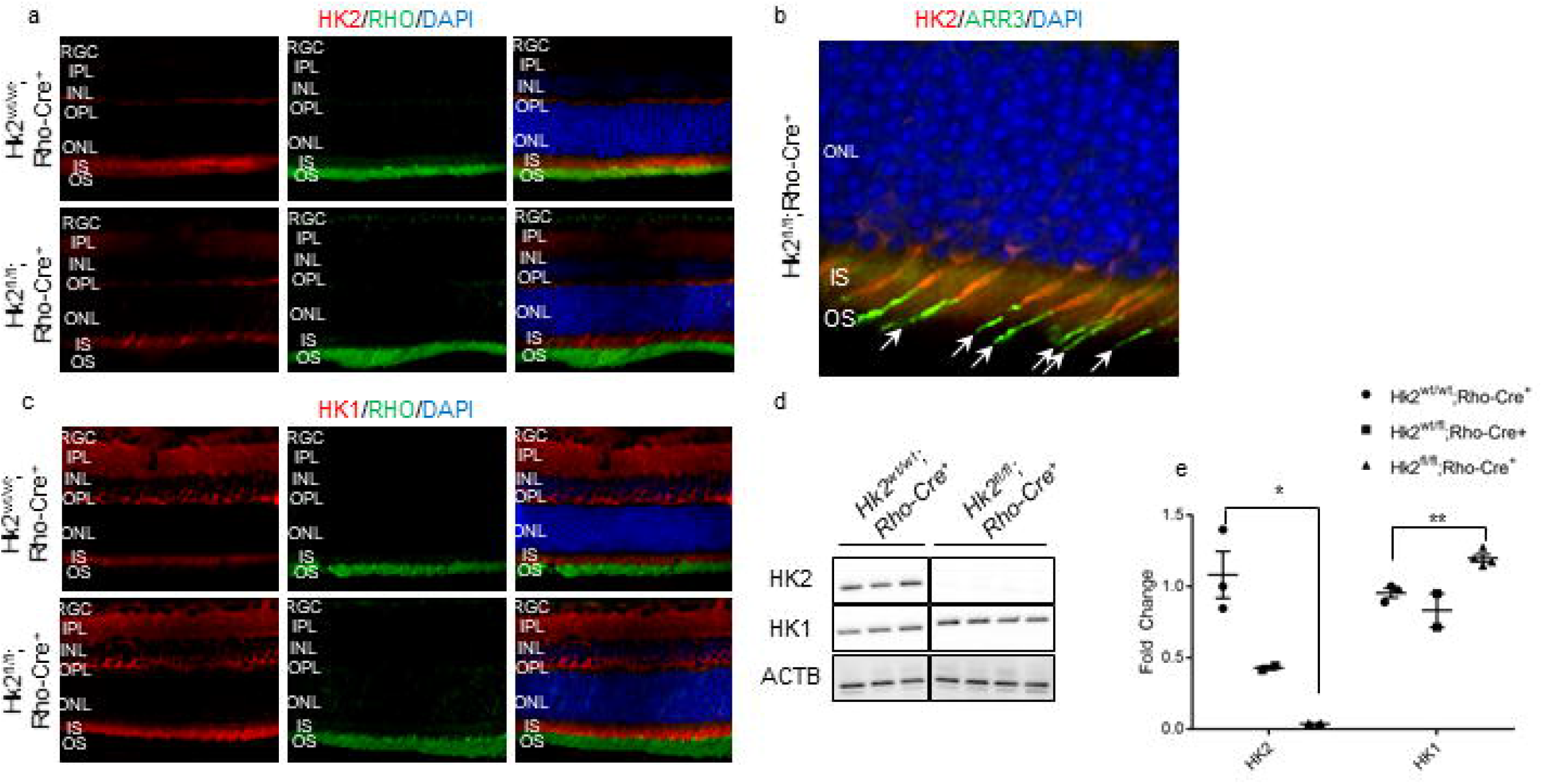
Successful knockdown of HK2 in rod photoreceptors with compensatory upregulation of HK1. (a) HK2 immunofluorescence (red) is found mainly in the inner segments of photoreceptors with little expression elsewhere in the retina of WT mice (*Hk2^wt/wt^;Rho-Cre*^+^). Rod photoreceptor-specific, *Hk2* cKO mice (*Hk2^fl/lt^;Rho-Cre*^+^) lack expression of HK2 in the majority of photoreceptors. (b) Co-labeling of HK2 (red) and ARR3 (green) confirms that the remaining HK2 expression is limited to cone photoreceptors in cKO mice. (c) HK1 immunofluorescence (red) depicts expression mainly in the inner retina of WT mice while cKO mice show upregulation of HK1 in photoreceptor inner segments. Nuclei of retinal cells are stained with DAPI (blue). (d) Western blot demonstrating almost complete loss of HK2 in cKO mice with a compensatory upregulation of HK1. ACTB (β-actin) was used as a loading control. (e) Quantitative analysis of HK2 and HK1 protein levels shows a statistically significant decrease in the level of HK2 in cKO mice. HK1 protein levels are statistically significantly increased in the retinas of these animals as compared to WT animals. N=3-4, *-p<.05 **-p<.01. RGC – Retinal Ganglion Cell layer, IPL – Inner Plexiform Layer, INL – Inner Nuclear Layer, OPL – Outer Plexiform Layer, ONL – Outer Nuclear Layer, IS – Inner Segments, OS – Outer Segments, DAPI – 4’ diamidino-2-phenylindole, RHO – Rhodopsin, ARR3 – Cone Arrestin

### HK2 promotes photoreceptor survival during acute outer retinal stress

As discussed above, one of the main non-enzymatic roles of HK2 is to inhibit apoptosis via its interaction with the mitochondria. The experimental model of retinal detachment utilized here results in increased apoptosis of photoreceptors.^28–33^ Furthermore, it was observed that HK2 was enriched in the mitochondria fraction after experimental retinal detachment in rats (Fig. 1d). Therefore, we sought to determine the role of HK2 in photoreceptor survival under acute outer retinal metabolic stress as produced by this experimental model of retinal detachment. The retinas of WT and cKO mice were detached and harvested 3 days later as this has been shown to be the peak of TUNEL staining in the outer nuclear layer (ONL) after retinal detachment and correlates with long-term photoreceptor survival.^7,34^ cKO mice had significantly more TUNEL positive cells in the ONL as compared to WT mice (Fig 3a&b). As in our rat model of retinal detachment (Fig. 1a), HK2 levels are significantly down regulated in WT mice 3 days post retinal detachment, and as expected, since cKO animals have significantly reduced levels of HK2 in the retina (Fig. 2), no change was detected following retinal detachment (Fig 3c). Surprisingly, HK1 protein levels were significantly down regulated in the retinas of cKO mice 3 days after retinal detachment, but not in the retinas of WT mice (Fig 3d).

**Figure 3.**
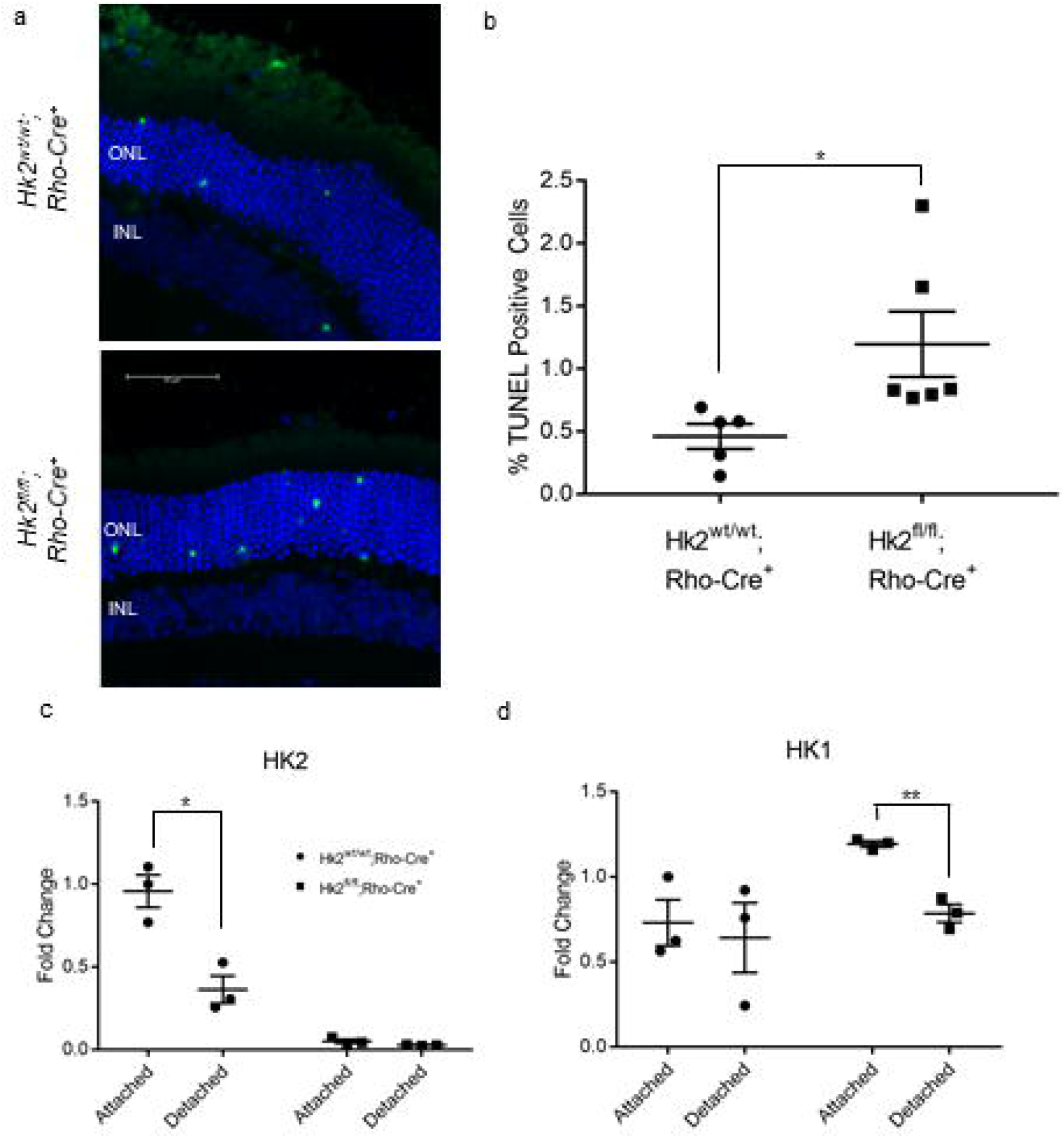
HK2 regulates photoreceptor survival after retinal detachment. (a) Representative images of TUNEL-stained photoreceptors (green) in detached regions of mouse retina 3 days post-retinal detachment. Nuclei of cells within ONL (outer nuclear layer) and INL (inner nuclear layer) are stained with DAPI (blue). (b) Quantification of TUNEL-positive cells in the ONL. N=5-6 eyes, p<0.05. (c) Quantitative analysis of HK2 protein levels 3 days after retinal detachment. HK2 is down-regulated following retinal detachment in WT mice. (d) Quantitative analysis of HK1 protein levels 3 days after retinal detachment. HK1 is downregulated following retinal detachment in cKO mice only. N=3-6 animals per group, *-p<0.05, **-p<0.001

#### HK2 affects photoreceptor survival and retinal function in aging

To assess the phenotype resulting from rod photoreceptor-specific, *Hk2* deletion, *in vivo* retinal morphology was assessed via optical coherence tomography (OCT). A small but significant decrease in OSEL (outer segment equivalent length) was observed at 1 month but was not present at 5 months of age (Fig. 4a&b). By 10 months of age, a significant decrease in total retinal thickness was observed in cKO mice as compared to WT mice (Fig. 4a&b). This thinning is driven by a reduction in ONL and OSEL thickness (Fig. 4b). Eyes from 10-month-old cKO and WT mice were sectioned and stained with hematoxylin and eosin to assess total ONL cell counts. The percent of ONL nuclei was statistically significantly less in cKO compared to WT mice (Fig. 4c&d), consistent with *in vivo* morphologic assessment. In accordance with these anatomic changes, visual function, as evaluated by electroretinography (ERG), showed no significant changes in any amplitudes at 2 and 5 months of age. Yet, at 10 months of age, statistically significant decreases were noted in the scotopic a-and b-wave amplitudes as well as in the photopic b-wave amplitude of cKO animals (Fig 4e-h).

**Figure 4.**
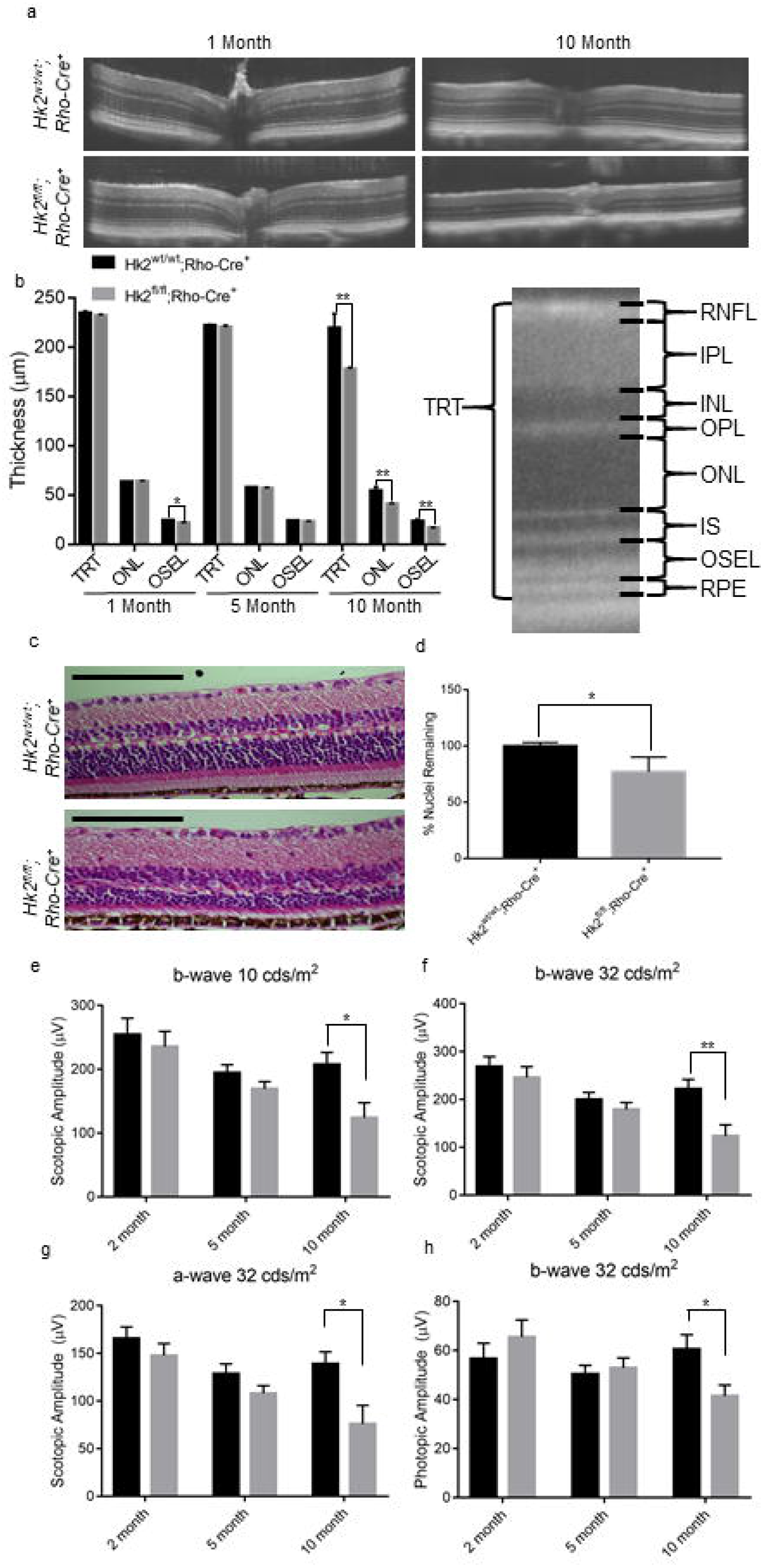
HK2 is required for long-term survival of photoreceptors. (a) Representative OCT images of 1- and 10-month-old cKO mice. (b) Quantitative analysis of retinal layers from OCT images. TRT, ONL and OSEL are significantly thinner in cKO mice as compared to WT animals. Schematic of retinal layers measured using OCT. OSEL is the region between the inner segment boundary and the apical surface of the RPE. (c) Representative hematoxylin and eosin stained sections of 10-month-old cKO and WT mice. (d) ONL nuclei counts normalized to total inner retinal area as a percent of the WT mice. (e-h) ERG amplitudes at indicated illuminance. (e-g) Scotopic and (h) photopic measurements. Amplitude is significantly decreased in cKO animals at 10 months of age. N = 8-16, * -p<.05, **-p<.01. TRT – Total Retinal Thickness, RNFL – Retinal Nerve Fiber Layer, IPL – Inner Plexiform Layer, INL-Inner Nuclear Layer, OPL – Outer Plexiform Layer, ONL – Outer Nuclear Layer, IS – Inner Segments, OSEL – Outer Segment Equivalent Length, RPE – Retinal Pigment Epithelium

### Deletion of HK2 does not significantly alter retinal metabolism

Rod photoreceptors have been reported to reprogram their energy metabolism from aerobic glycolysis to oxidative phosphorylation when *Hk2* is deleted.^26^ To investigate this potential metabolic reprogramming in our rod-specific *Hk2* cKO model, the transcriptional profile of central glucose metabolism genes was determined in the retina. We conducted real-time PCR using the Mouse Glucose Metabolism RT^2^ Profiler™ PCR array (Qiagen), which assesses genes involved in glycolysis, gluconeogenesis, tricarboxylic acid cycle, pentose phosphate pathway, glycogen synthesis, glycogen degradation, and regulation of glucose and glycogen metabolism. Of the 84 genes examined, only seven showed significant (p<0.05) expression differences between the *Hk2* cKO mice and WT mice (supplemental table 1, Fig. 5a). Of these, only *Fbp2* demonstrated a statistically significant fold change greater than 2. FBP2 (fructose 1,6-bisphosphatase, 2.9 fold change) is the rate-limiting step of gluconeogenesis and catalyzes the conversion of fructose 1,6-bisphosphate to fructose 6-phosphate. Conversion of glucose to lactate is a hallmark of aerobic glycolysis. Photoreceptors process 80-96% of all glucose into lactate; therefore, a decrease in the total steady-state lactate levels would suggest significant perturbations to aerobic glycolysis.^35^ No significant differences in total retinal lactate levels between cKO and WT animals were observed at 2 months of age (Fig. 5b). Similarly, PKM2 and LDHA, the two other main regulators of aerobic glycolysis, were assayed by Western blotting. No significant differences in the expression levels of either enzyme was detected between cKO and WT mice (Fig. 5c-f). Collectively, these data demonstrate the lack of any major retinal metabolic reprogramming of glucose metabolism when *Hk2* is genetically deleted in rod photoreceptors.

**Figure 5.**
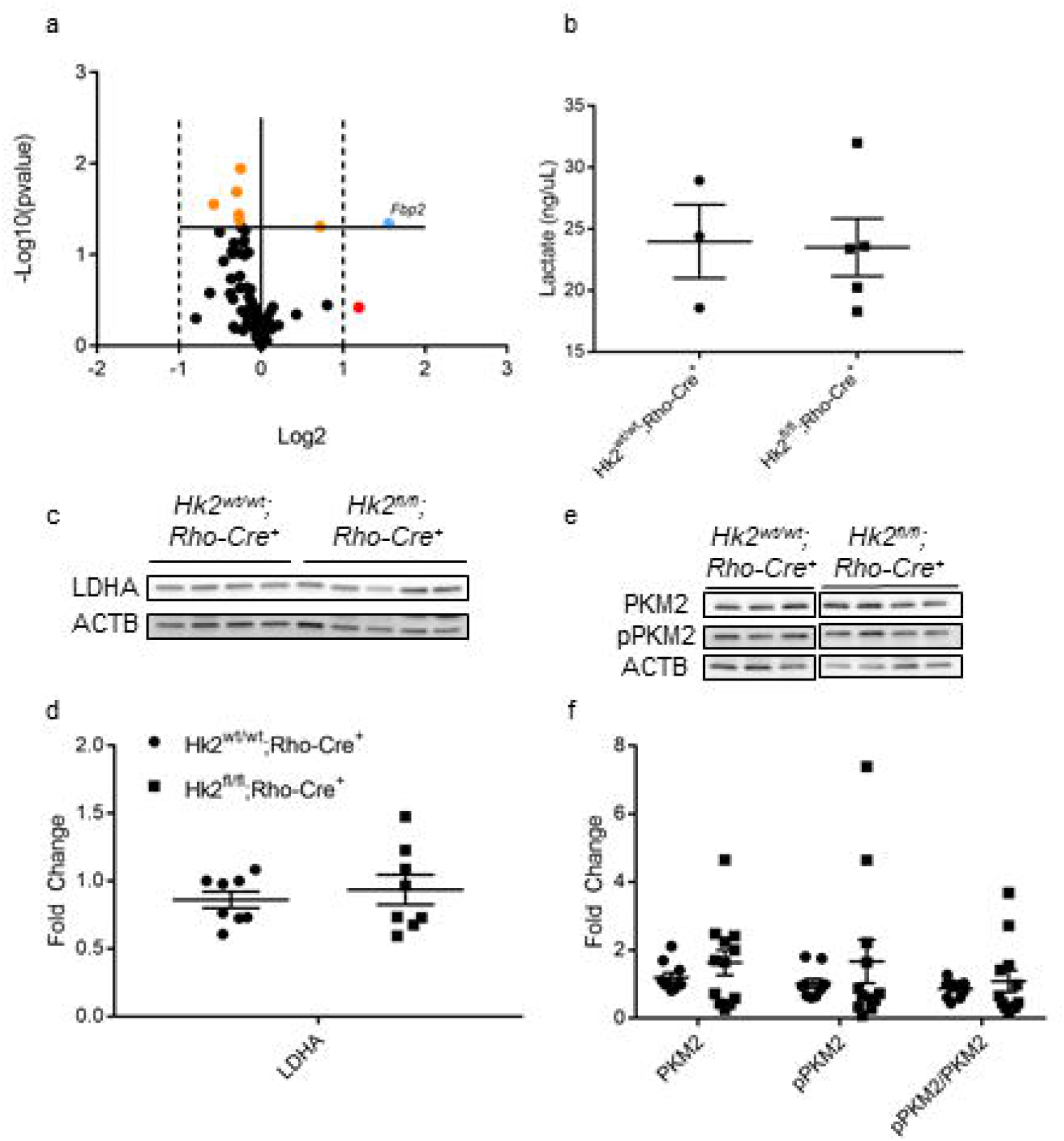
Glucose metabolism is largely unchanged in rod photoreceptor-specific, *Hk2* conditional knockout mice. (a) Volcano plot of 84 glucose metabolism gene expression as measured by qRT-PCR. The solid horizontal line indicates a p-value of 0.05. The vertical dotted lines indicate a fold change of ± 2. Orange dots are significantly different. The red dot has an insignificant fold change greater than 2. The blue dot shows a gene changed more than ± 2 fold and with a p-value of <0.05. Only *Fbp2* is significantly changed greater than 2-fold. (b) Total retinal lactate is unchanged between cKO and WT mice measured at 6-8 weeks of age. (c&d) Western blot and respective quantitative analysis confirms LDHA protein levels are unchanged after deletion of HK2 from rod photoreceptors. (e&f) Western blot of total PKM2 and p-PKM2 (Y105) and respective quantitative analyses demonstrates that these levels are unchanged in cKO animals as compared to WT week old mice. N = 3-9, mean + SEM.

## Discussion

In this study, we show that HK2 translocates to mitochondria following retinal detachment and at the same time is significantly downregulated. This subcellular localization following nutrient deprivation stress in the retina may be mediated by increased HK2 phosphorylation via activated retinal AKT. We also show that inhibiting AKT phosphorylation using a broad PI3K inhibitor (LY294002) results in accumulation of HK2 in the cytosol. Treatment with LY294002 activates caspase 3, 7, and 8, and reduces cell viability, possibly due to reduced HK2 fraction in the mitochondria. Rod-specific *Hk2* deletion results in the upregulation of *Hk1* expression in the photoreceptor inner segments. Under acute outer retinal stress induced by experimental retinal detachment, HK2-to-HK1 isoform switching increases rod photoreceptor susceptibility to cell death. This HK isoform switch also leads to degenerative changes in the outer retina with aging, as shown by the decrease in both ERG amplitudes and concurrent decline in retinal thickness measurements. Interestingly, only a single central metabolic gene, *Fbp2*, exhibited a significant change in expression and no significant changes in aerobic glycolysis were observed in the retinas of the rod photoreceptor-specific, *Hk2* cKO mice. These data suggest that increased HK1 expression in rods functionally replaces HK2 at the metabolic level. Under the experimental conditions used in these studies, the neuroprotective effect of HK2 during acute outer retinal stress and aging may be due to its non-enzymatic activities.^15,16,18^

Deletion of HK2 from rod-photoreceptors led to an increase in total HK1 levels in the retina, specifically in the rod photoreceptor inner segments. Despite HK2 expression being associated with increased aerobic glycolysis, we were unable to detect any significant perturbations to aerobic glycolysis in photoreceptors lacking HK2.^36,37^ For example, lactate is almost exclusively produced by photoreceptors in the retina through aerobic glycolysis, however no significant changes in steady-state lactate production were observed in the *Hk2* cKO retinas.^35^ Furthermore, very few changes were observed in the expression of genes involved in central glucose metabolism. Taken together, these data suggest that HK1 can metabolically replace HK2. It will be important to both determine the steady-state levels of the metabolites during glucose metabolism as well as the flux of ^13^C-glucose as it is broken down through glycolysis and branching pathways to examine the exact metabolic effects of HK2-to-HK1 substitution in photoreceptors.

As glucose is the primary fuel source for photoreceptors, an isoform of hexokinase is likely required to initiate the glycolytic cascade. All isoforms of hexokinase perform the same enzymatic reaction, phosphorylating glucose to glucose-6-phosphate. Yet, each isoform has differing enzymatic rates to account for varying responsibilities depending on cell type.^38^ Interestingly, HK1 and HK2 have similar enzymatic constants and even though HK2 is believed to be critical for aerobic glycolysis, this enzymatic similarity may allow HK1 to maintain metabolic homeostasis and ultimately, to prevent significant transcriptional response to compensate for the absence of HK2. The only gene which was significantly altered in *Hk2* cKO retinas was *Fbp2.* Enhanced fructose bisphosphatase (FBPase) activity and decreased fructose 2,6-bisphosphate (F2,6P) levels secondary to increased *Fbp2* expression would be expected to decrease glycolytic flux and may maintain photoreceptor glucose homeostasis in *Hk2* cKO retinas via opposing PFK1 activity, the critical enzyme for committing glucose-derived metabolites to glycolytic flux.^39,40^

The data presented here suggest that the non-enzymatic functions of HK2 may be more important for preserving photoreceptors during acute stress than its enzymatic roles. We found that the distribution of HK2 is nearly equal between cytosolic and mitochondrial fractions in the normal rat retina. It has been shown that both HK1 and HK2 are maximally enzymatically active and preferentially use mitochondrial produced ATP to phosphorylate glucose.^41^ Since not all of HK2 is found in the mitochondrial fraction, HK2 may be regulated to fine tune the amount of glucose entering glycolysis. Alternatively, HK2 may be performing a critical, non-enzymatic function in the cytosol unrelated to glycolysis. Further studies are needed to investigate the potentially distinct roles of HK2 in the cytosol, which may explain why HK2 is preferentially expressed in photoreceptors. One of the major non-enzymatic roles of HK2 is to inhibit binding of pro-apoptotic factors and prevent opening of the mPTP, which may explain why HK2 translocates to the mitochondrial fraction following retinal detachment. Experimental retinal detachment induces acute apoptotic stress on photoreceptors mediated by the Fas signaling cascade.^29,33^ When HK2 translocation is partially blocked using the PI3K inhibitor LY294002, 661W cells activate caspases and undergo cell death. While inhibiting the PI3K signaling cascade alters numerous intracellular pathways that may be important in cell survival, our data suggests that HK2 binding to mitochondria is at least partially responsible for inhibiting apoptosis, similar to what was shown in other experimental models.^19^

Photoreceptors lacking HK2 are more susceptible to acute nutrient deprivation caused by retinal detachment. Since HK1 replaces HK2 in cKO photoreceptors and has a similar affinity for glucose when compared to HK2, this susceptibility to nutrient deprivation is likely not related to metabolizing the limited amount of glucose in the detached retina.^41^ Interestingly, HK2 levels drop precipitously 3 days after retinal detachment in WT animals. Likewise, when HK2 is replaced by HK1 in *Hk2* cKO retinas, a robust decrease of HK1 expression was also observed after retinal detachment, in contrast to stable HK1 levels in WT retinas. These findings indicate that HK levels are regulated in rod photoreceptors during stress. The exact mechanisms of how HK2 production or degradation is regulated need to be explored further to fully understand its role for neuroprotection. Regardless of what is precipitating HK2 loss, it may be an attempt to catabolize intracellular proteins through autophagy to produce additional energy substrates and prolong survival in a nutrient deficient state. The degradation of HK2 during metabolic stress may disturb the intricate balance of apoptotic inhibition and energy production, uncoupling metabolic stress from cell death signaling and increasing the sensitivity of photoreceptors to nutrient deprivation. Raising the overall levels or preventing the degradation of HK2 may be beneficial for boosting photoreceptor survival during nutrient deprivation.

Even though HK1 appears to functionally replace HK2 at the metabolic level, *Hk2* cKO retinas still developed an age-related phenotype. Similar to a previous report, we observed rod dysfunction in older *Hk2* cKO mice as measured by ERG.^26^ The functional phenotype identified by Petit and colleagues was not associated with any morphological change in retinal structure. In contrast, we show that rod ERG dysfunction was associated with significant outer retinal abnormalities anatomically. This difference may be due to variations in genetic background between the two *Hk2* cKO strains. We screened our animals for the *rd8* mutation to rule out the possibility of background degeneration and found that our *Hk2* cKO animals lack the *rd8* allele. In addition, the phenotype of the *rd8* mutation has a very rapid onset, around 6 weeks of age, and is characterized by focal retinal folds, pseudorosettes, and retinal degeneration.^42^ The phenotype observed in the HK2 cKO mice here was a slowly progressing degeneration and is not characteristic of the *rd8* mutation or other common background degenerative retinal diseases in mice.^42^

Our results suggest that HK2 has an essential function in preserving photoreceptors during aging. The time points we examined demonstrate that the onset of retinal degeneration in HK2 cKO animals is delayed. Our functional analysis by ERG and *in vivo* anatomic analysis by OCT at 5 months of age show that the overall retinal health is normal at this time. Thus, any potential degeneration was below the threshold of detection or had not begun yet. Yet, by 10 months of age there was a dramatic decrease in ONL and OSEL thickness. Interestingly, others have shown that the transcriptional profile of rod photoreceptors begins to change by 5 months of age.^43^ Parapuram and colleagues found differential regulation of genes belonging to many different signaling cascades, including apoptosis, as animals age. At the same time, photoreceptors begin to accumulate significant oxidative damage with age, which may become toxic and increase apoptotic signaling.^44^ The combination of a loss of HK2, changes in gene regulation, and the accumulation of oxidative injury may result in an imbalance in apoptotic signaling leading to delayed photoreceptor degeneration in our mouse model. Further studies are needed to determine the time course of photoreceptor degeneration in the *Hk2* cKO animals with age and investigate the enzymatic and non-enzymatic functions of HK2 in aging photoreceptor cells.

Based on our data, we propose the following model for HK2 function in photoreceptors (Fig. 6). During nutrient homeostasis, HK2 is equally distributed between the cytosolic and mitochondrial compartments. After retinal detachment, glucose levels drop precipitously. This results in a redistribution of HK2 to the mitochondrial fraction. At the same time, AKT is activated, phosphorylating HK2 and enhancing its binding to VDAC. This then serves to prevent the binding of pro-apoptotic factors to the mitochondria and prevent apoptotic signaling. Since the total amount of HK2 drops significantly after retinal detachment, the neuroprotective effect of HK2 mediated by mitochondrial regulation of apoptosis may be limited. Determining the interactions between AKT/VDAC/HK2 in the retina will be important for verifying this model. These experiments will enhance our understanding of the role of HK2 in both anti-apoptotic and pro-survival functions in photoreceptors.

**Figure 6.**
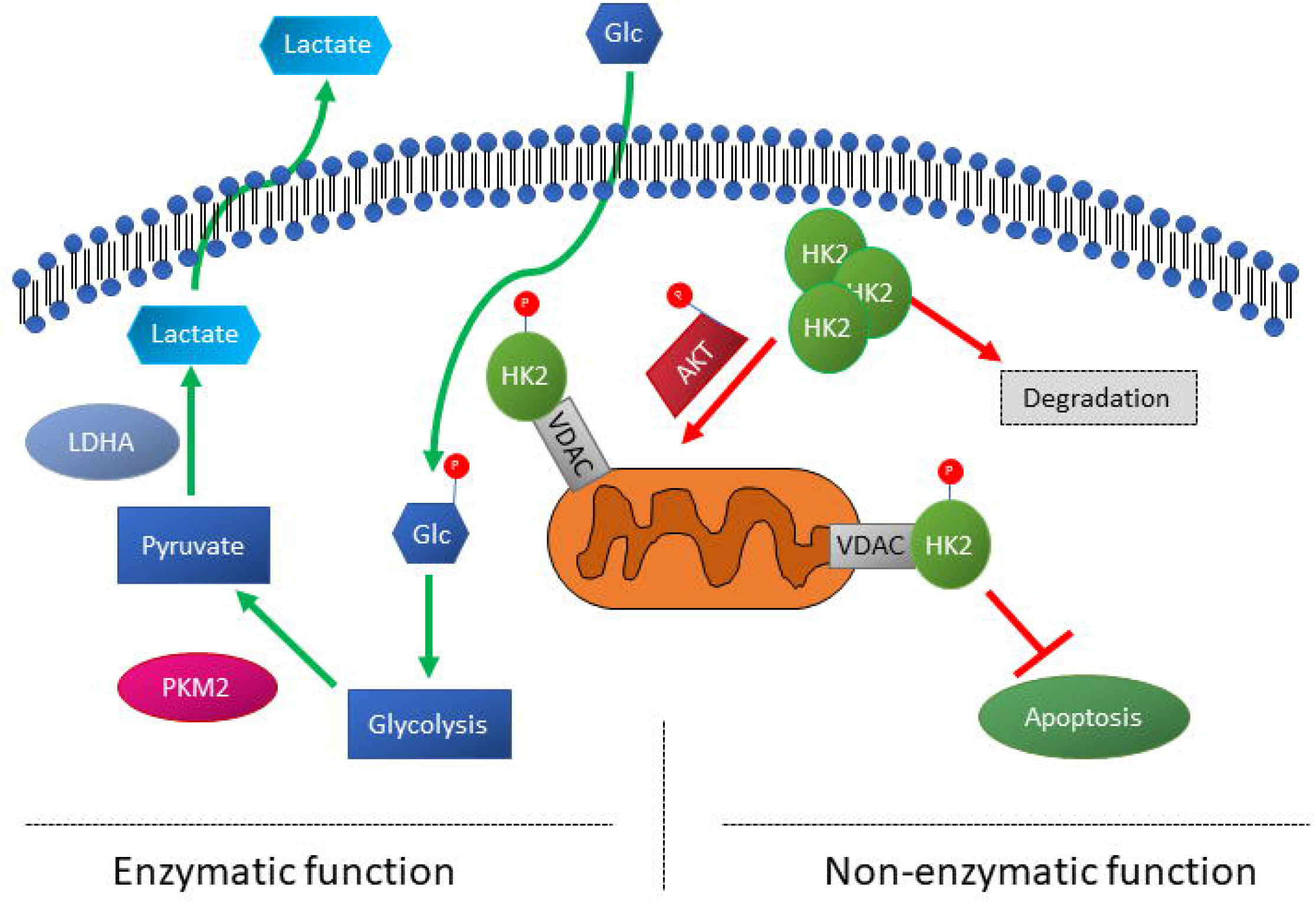
Proposed model of HK2 activity during nutrient stress.

We previously showed that reprogramming photoreceptor metabolism via modulating a key regulatory enzyme of aerobic glycolysis, PKM2, is a novel therapeutic strategy. Data presented in this study demonstrate that modulating the other critical regulator of aerobic glycolysis, HK2, may also increase photoreceptor survival during metabolic stress, potentially secondary to its non-enzymatic functions. Determining the exact role of HK2 during nutrient stress has the potential to reveal novel molecular pathways that can be targeted for photoreceptor neuroprotection.

## Methods

### Animals

All animals were treated in accordance with the Association for Research in Vision and Ophthalmology (ARVO) Statement for the Use of Animals in Ophthalmic and Vision Research. The protocol was approved by the University Committee on Use and Care of Animals of the University of Michigan (Protocol number: PRO00007463). All animals were maintained at room temperature in a 12/12-hour light/dark cycle. Mice harboring a floxed *Hk2* gene were a gift from Dr. Mohanish Deshmukh and Dr. Timothy Gershon.^25^ These mice were crossed to mice carrying a Cre-recombinase gene under the control of the rhodopsin promoter.^24^ The mice were maintained on a C57BL/6 background and were confirmed to not carry the *rd8* mutation. *Hk2^wt/wt^;Rho-Cre^+^*(Wildtype, WT) *and Hk2*^*fl/fl*^;*Rho-Cre*^+^ (conditional knockout, cKO) mice were generated.

### Sub-cellular Fractionation

Rodent retinas were isolated from euthanized animals using the “Cut and Pick” (or Winkling) method while being careful to avoid collecting RPE.^45^ The retinas were fractionated into cytosolic and post-cytosolic fractions using the Subcellular Protein Fractionation Kit for Tissues (Thermo Fisher Scientific; Waltham, MA, USA; Cat# 87790) following a modified manufacturer’s protocol. Two retinas were pooled for each sample and homogenized in cytoplasmic extraction buffer (CEB) supplemented with protease (Halt™ Protease Inhibitor Cocktail, Thermo Fisher Scientific, Cat# 87787) and phosphatase (Halt™ Phosphatase Inhibitor Cocktail, Thermo Fisher Scientific, Cat# 78420) inhibitors using 30-40 strokes with the “B” pestle of a Dounce homogenizer. Lysate was then centrifuged at 4°C for 10 minutes at 10,000 relative centrifugal force (RCF). The supernatant (cytosolic fraction) was saved and the pellet was sonicated at 20% amplitude with 1 second on/off pulse for 10 seconds in RIPA lysis buffer (Thermo Fisher Scientific, Cat# 89900) supplemented with protease and phosphatase inhibitors. This lysate was then centrifuged at 4°C for 10 minutes at 10,000RCF. The supernatant (mitochondrial enriched fraction) was saved and the pellet was discarded.

The 661◻W photoreceptor cell line was a generous gift from Dr. Muayyad Al-Ubaidi.^46^ 2×10^6^ 661W cells were seeded 24 hours prior to treatment. Cells were fractionated using the Subcellular Protein Fractionation Kit for Cultured Cells (Thermo Fisher Scientific, Cat# 78840) as above except cells were trypsinized, pelleted, rinsed with 1X phosphate buffered saline (PBS), re-pelleted before being resuspended in CEB for a 10-minute incubation at 4°C. The remaining steps were identical. Cells were cultured as described previously and treated with DMSO or LY294002 (Cell Signaling Technology; Denver, MA, USA, Cat# 9901, 150μM, 5mL) in glucose-free DMEM (Thermo Fisher Scientific; Cat #11966025) supplemented with 5.5mM Glucose for 1.5 hours prior to fractionation.^47^ Whole cell lysates were also collected to verify loss of p-AKT after 1.5 hours of LY294002 treatment.

### Caspase Activation and Cell Viability Assays

661W cells were seeded at a density of 10,000 cells/well in each well of a 96-well plate then treated 24 hours later with DMSO, 50μM LY294002, or media alone for 6 hours. Caspase activity was assayed using the Caspase-Glo® 3/7 or 8 Assay System (Promega; Madison, WI, USA, Cat# G8090 or G8200) and cell viability was measured using the CellTiter-Glo® Luminescent Cell Viability Assay (Promega, Cat#G7570) following the manufacturer’s supplied instructions. Briefly, cells were treated for the desired length of time and then prepared reagent was added directly to each assay well, including blank wells containing media alone or media containing either DMSO or LY294002. The reaction was allowed to incubate for 15 minutes before assaying using a luminometer.

### Western Blotting

After harvest, retinas were homogenized in RIPA lysis buffer supplemented with protease and phosphatase inhibitors. Homogenization was performed using a sonicator set at 20% amplitude with 1 second on/off pulse for 10 seconds followed by centrifugation at 10,000RCF at 4°C for 10 minutes. Clarified lysates had protein content estimated using the Pierce™ BCA Protein Assay Kit (Thermo Fisher Scientific, Cat# 23225). 15μg of total protein was diluted in Laemmli buffer (Bio-Rad; Hercules, CA USA, Cat# 1610747) supplemented with β-mercaptoethanol (Millipore-Sigma; St. Louis, MO USA, Cat# M6250). Denatured protein samples were run on 4–20% or 10% Mini-PROTEAN® TGX™ Precast Protein Gel (Bio-Rad, Cat# 4561094). Protein was transferred using either a wet-transfer (100V for 1 hour at 4°C) or the Trans-Blot® Turbo™ Transfer System (25V for 30 minutes) (Bio-Rad, Cat# 1704150). After transfer, the PVDF membranes were blocked using 5% non-fat dry milk in TBS-T (Tris-buffered Saline (Bio-Rad, Cat# 1706435) supplemented with 0.1% Tween-20 (Thermo Fisher Scientific, Cat# 28320)). Primary antibodies were diluted according to Table 1 in 5% Bovine Serum Albumin (BSA; (Millipore-Sigma, Cat# A9647)) in TBS-T and blocked membranes were incubated overnight at 4°C with gentle agitation. Membranes were then washed and incubated with appropriate secondary antibody diluted in 5% BSA in TBST for 1 hour at room temperature. Protein bands were detected using the SuperSignal™ West Dura/Femto Extended Duration Substrate (Thermo Fisher Scientific, Cat# 34075 and 34094) with an Azure c500 imaging system (Azure Biosystems; Dublin, CA USA). Densitometry was performed using the Azure Spot program (Azure Biosystems; Dublin, CA USA).

**Table 1.**
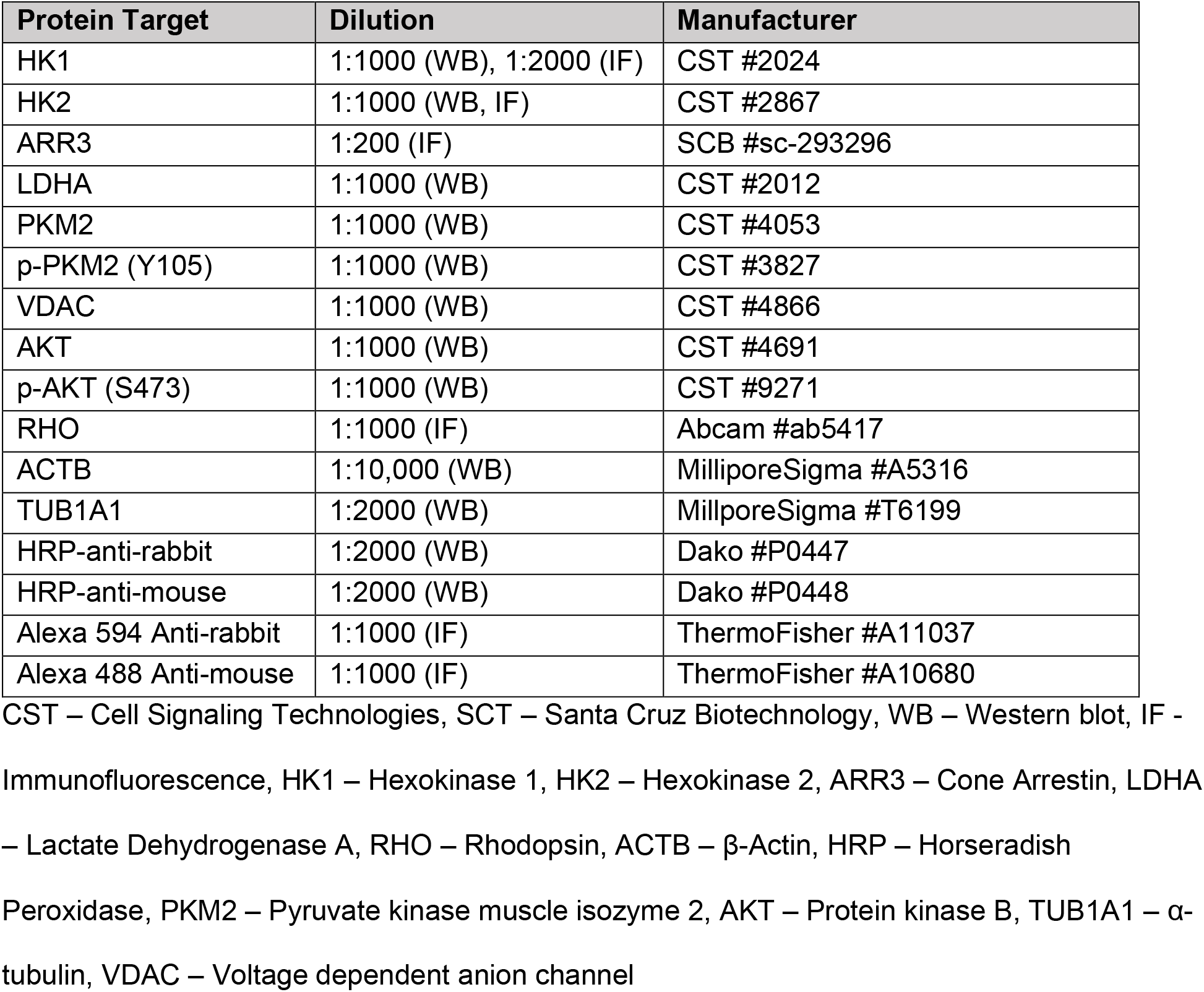
Antibodies used

### Immunofluorescence

Mouse eyes were enucleated and fixed in 4% formaldehyde (Polysciences; Warrington, PA, USA, Cat# 18814) overnight at 4°C. Fixed eyes were embedded in paraffin and sectioned 6μm thick. Sections were de-paraffinized and antigen retrieval was performed following standard procedures. Blocking was performed with 1% BSA in PBST supplemented with 10% Normal Goat Serum (NGS) for 1 hour before adding primary antibody (Table 1) diluted in 1% BSA supplemented with 1% NGS overnight at 4°C. Slides were then washed in PBS and then with 1% BSA in PBST supplemented with 1% NGS before appropriate secondary antibody was added (Table 1) and slides were incubated at room temperature for 1 hour. Finally, slides were washed with PBS and counterstained with ProLong™ Gold Antifade Mountant with DAPI (Thermo Fisher Scientific, Cat# P36930). Slides were imaged using a Leica DM6000 microscope with a 40X objective.

### qRT-PCR

Retinas were extracted as described above and immediately immersed in RNAlater (Qiagen; Hilden, Germany, Cat# 76104). Total RNA was extracted using the RNeasy Mini Kit (Qiagen, Cat# 74104) following the manufacturer’s instructions. RNA quantity and quality were assessed with a Nanodrop 1000 (Thermo Fisher Scientific). 400 ng of total RNA was reverse transcribed using the RT^2^ First Strand Kit (Qiagen, Cat# 330401). Transcriptional changes in metabolic pathways was assessed using the RT^2^ Profiler PCR Array for Mouse Glucose Metabolism (Qiagen, Cat# 330231 PAMM-006ZA) following the manufacturer’s instructions. 102uL of cDNA was pre-mixed with 650uL of 2X SYBR Green master mix (Qiagen, Cat# 330503) and 548uL of ddH2O. 10uL of each pre-mix was added to each well of the 384 well plate following the manufacturer’s instructions. The plates were thermocycled using a CFX384 thermocycler (Bio-Rad) following the provided cycling parameters. The geometric mean of the Ct values for *Actb*, *Gapdh*, and *Hsp90ab1* was used for relative quantitation using the 2^-ΔΔCt method.

### Functional Assessment

Functional assessment of mouse vision was performed as described previously.^7^ Total retinal function was measured using electroretinography (ERG). Mice were anesthetized and the ERG response was measured using the Diagnosys Celeris ERG system (Diagnosys LLC, Lowell, MA, USA). *In vivo* retinal thickness was assessed using the Envisu-R SD-OCT imager (Leica Microsystems Inc., Buffalo Grove, IL, USA). Average thickness of various retinal layers was assessed using Diver 1.0 (Leica Microsystems Inc). Four points 200μm from the optic nerve head were used to measure layer thickness and the results were averaged. Total retinal thickness (TRT), outer nuclear layer (ONL) thickness and outer segment equivalent length (OSEL) were determined. OSEL is defined as the length from the ends of the inner segment boundary to the surface of the RPE.^7,48^

### Retinal Detachment, TUNEL Staining, and ONL Counts

Experimental neurosensory retinal detachment was induced in mice as described previously.^7^ Briefly, animals were anesthetized and a sclerotomy was performed using a 25G micro-vitrealretinal blade. A 35G beveled cannula (World Precision Instruments; Sarasota, FL, USA, Cat# NF35BV-2), attached to a NanoFil-100 syringe (World Precision Instruments, Cat# NANOFIL-100) was used to inject 2-3μL of Healon (1% Hyaluronic Acid) (Abbott Medical Optics; Santa Ana, CA, USA, Cat# 05047450842) between the photoreceptor and RPE layers. Care was taken to detach approximately half of the retina in each animal. Only the sclerotomy was performed on the fellow eye as a control. After 3 days, animals were sacrificed and whole eyes were extracted and embedded in paraffin for sectioning as described above. TUNEL staining was performed using the DeadEnd™ Fluorometric TUNEL System (Promega Cat# G3250) and sections were counterstained using ProLong Gold Mountant with DAPI. Stained sections were imaged with a Leica DM6000 microscope using a 40X objective. TUNEL positive cells were manually counted across the detached portion of the retina. Counts were normalized to the total number of nuclei in the outer nuclear layer in the detached region, counted manually or with an automated cell counting macro using ImageJ.^7,47^

### Total Retinal Lactate Quantification

Total Retinal Lactate was quantified using the Lactate Assay Kit (Millipore-Sigma, Cat #MAK064) following the manufacturer’s supplied protocol. Animals were euthanized and neural retina was extracted as described above and briefly washed in PBS to remove any adhered vitreous. The retinas from both eyes of each animal were pooled and homogenized using the provided Lactate Assay Buffer. A sample of homogenate was retained for total protein quantification to normalize samples before de-proteinization using the Amicon Ultra-0.5 Centrifugal Filter Unit (Millipore-Sigma, Cat # UFC501024), which has a molecular weight cut-off of 10kDa. 10μL of sample was loaded in duplicate for both the colorimetric and fluorometric assay. A standard lactate concentration curve was plotted for each assay and the concentration of each sample was determined from this curve. The concentration of lactate (ng/μL) was normalized by the total sample protein concentration.

### Statistical Analysis

Statistical analysis was performed comparing WT to cKO samples using a two-tailed student’s t-test with either Excel or Prism 7.0. Results with a p-value ≤ 0.05 were considered significant. Data are displayed as mean ± SEM, and the number of replicates is indicated in each figure legend.

## Acknowledgements

The authors would like to acknowledge Sarah Sheskey for her help performing ERG. This work was supported with a grant from the Macula Society, E. Matilda Ziegler Foundation, and from the NEI (5K08EY023982). This work utilized the Vision Research Core funded by P30 EY007003 from the National Eye Institute.

## Data Availability

All data will be provided upon reasonable request to the corresponding author.

